# Integrated detection and quantification of aberrant transcripts with novel splicing events

**DOI:** 10.1101/2025.02.02.635191

**Authors:** Ira Agrawal, Yong Shan Lim, Emma May Sanford, Wan Yun Ho, Shuo-Chien Ling, Greg Tucker-Kellogg

## Abstract

Splicing misregulation, such as the inclusion of previously unknown cryptic exons, is implicated in numerous diseases. Recent methods have increased accurate and efficient detection of such splicing alterations occurring in disease phenotypes. However, the quantification and differential analyses of non-canonical splicing alterations remains focused at a splice event level, thus preventing a complete view of the effects on the downstream transcriptomic landscape. Here, we present a novel and integrated pipeline, SpliCeAT, that (1) detects and quantifies differential non-canonical splicing events from short-read bulk RNA-seq data, (2) augments the canonical transcriptome with novel transcripts containing these non-canonical splicing events, and (3) performs transcript-level differential analysis to identify and quantify aberrant cryptic exon-containing transcripts based on this *augmented transcriptome*. Using TDP-43, an ALS/FTD-associated RNA-binding protein as an example, we identified and catalogued aberrant splicing events in embryonic mouse brains from deletion of TDP-43 in neural progenitor cells. The accuracy of our integrated pipeline was further confirmed and validated with long-read isoform sequencing. Furthermore, by comparing neuronal TDP-43 knockouts in mice with a publicly available human dataset with TDP-43 pathology, we identified and validated 4 common genes, namely, Kalrn/KALRN, Poldip3/POLDIP3, Rnf144a/RNF144A, and Unc13a/UNC13A, with cryptic exons. In summary, our integrated pipeline, novel splice events are identified, incorporated and quantified at the transcript level, thereby enabling more complete transcriptome profiling of well-annotated genomes in in the case of pathological splicing misregulation.

## Background

Almost all mRNAs in eukaryotes are products of splicing, in which a single pre-mRNA can yield multiple alternatively spliced mature transcripts. Precise regulation of alternative splicing is essential for maintaining the integrity and diversity of functional eukaryotic transcripts and their downstream translation products. More specifically, transcript diversity through alternative splicing is especially prevalent in tissues of the central nervous system [1–3]. Gene annotation systems and large scale studies try to capture the full range of non-pathological transcript diversity [4, 5].

Aberrant splicing produces undesired splicing products, some of which include cryptic exon events — e.g. cassette exons, exon extensions, and intron retentions — and others which involve exon skipping. These aberrant splicing products may be targeted for nonsense-mediated decay in the cytoplasm or be translated to dysfunctional proteins. Splicing misregulation is implicated in numerous diseases, such as amyotrophic lateral sclerosis (ALS) [6–12], Alzheimer’s disease [13, 14] and cancer [15]. In ALS, the functional lack of the RNA-binding protein TDP-43 in the central nervous system leads to inclusions of cryptic exons in a multitude of transcripts, of which cryptic splicing of Stathmin-2 and UNC13A have been shown to be disease-causing [6–12].

A wide range of approaches have been developed for detecting novel splicing events from short-read bulk RNA-seq datasets [15–25]. However, the tools developed for these approaches often report varying outputs, focus on distinct types of splicing events, or present results at different levels of granularity. When they can be compared, the results between the tools show little overlap [26], with a large number of reported false positive events. There is thus a need for an integrative approach that summarises detected splicing event output from multiple tools and consolidates them into a list of high-confidence, non-redundant novel splicing events. In line with this, Fenn *et al* [26] have recently developed a pipeline which integrates eight splicing event detection tools and produces a uniform report unifying their respective splicing output formats. Such integrative approaches are imperative in preventing over-reliance on a specific tool and avoiding tool-specific inherent biases.

Novel splicing event detection tools have focused primarily on identifying local splicing events, rather than characterising the transcripts from which they arise. For a more comprehensive understanding of how novel splicing affects the transcriptome landscape, there is a need to depart from the traditional single splice event-level analysis, and move towards transcript-level analysis. This involves studying how novel splice events are included into novel transcripts and subsequently how the expression of these transcripts differs between samples.

In this study, we seek to take full advantage of reference annotation for differential transcriptomics while working in pathological settings characterised by splicing misregulation. We address these issues by developing a novel pipeline, SpliCeAT, which integrates various splice event detection tools to (1) detect high-confidence novel splicing events, (2) create an augmented transcriptome to capture the inclusion of these high-confidence novel splicing events at the transcript level, and (3) use the augmented transcriptome for accurate quantification of global isoform composition changes. In order to evaluate and illustrate our pipeline, we used short-read bulk RNA sequencing data from brain tissues of 14.5-day embryonic mice with TDP-43 deleted from neural progenitor cells. We validate the isoforms and novel splicing events using long-read isoform sequencing (Iso-Seq).

## Results

### Overview of SpliCeAT’s differential splicing detection framework

To provide a more complete view of the effects of differential splicing on the downstream transcriptomic landscape, we developed an integrated pipeline (SpliCeAT) for quantification and differential expression analysis of transcripts containing novel events. Figure 1a provides an overview of the pipeline framework, starting from RNA-seq read alignment and transcript assembly, followed by differential splicing detection and generation of the augmented transcriptome.

**Figure 1.**
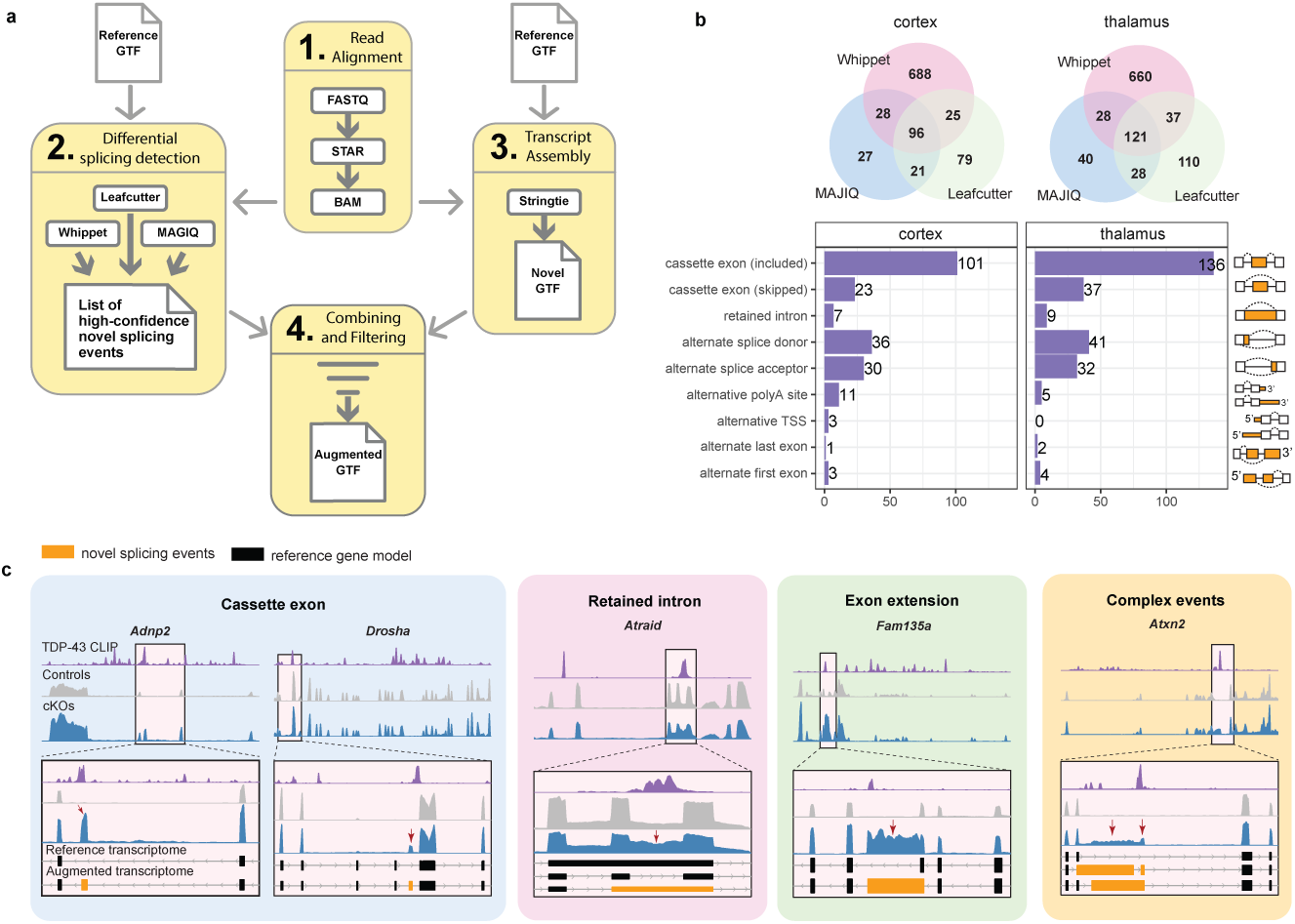
SpliCeAT framework and high-confidence differential splicing events. (a) SpliCeAT framework (b) Differential splicing events identified in cortex and thalamus tissue of TDP-43 cKO mice versus control mice via Whippet, MAJIQ and Leafcutter, along with the distribution of the types of high confidence novel splicing events detected. Venn diagrams showcase the gene-wise overlap between tools, while bar graphs showcase the number of the different types of differential splicing events. Note that there can be more than one differential splicing event occurring in a gene. (c) Visualisation of various splicing events: cassette exons, retained intron, exon extension and complex events. Upper panels show the entire gene region with tracks from TDP-43 CLIP-seq binding sites andshort-read RNA-Seq reads from control and cKO mice. The lower panel is zoomed in to the regions containing novel splicing events with the same tracks in addition to gene models.

After aligning short-read bulk RNA-seq reads using a splice-aware aligner, the next step of SpliCeAT is to detect high-confidence differential splicing events. A recent study comparing eleven alternative splicing event detection tools using simulated data [26] reported that MAJIQ [18] and Whippet [17] performed best for detecting *de novo*, unannotated events. Additionally, Whippet accurately detected the highest proportion of alternative splicing events. Thus, we used both MAJIQ and Whippet, together with another widely used tool, Leafcutter [16], to detect alternative splicing events in our RNA-seq reads. The three tools use different approaches to detect novel splicing events. MAJIQ first builds a splice graph for each gene and subsequently detects all local splicing variations (LSVs) present in a gene, and then tests for differential LSVs between conditions. Whippet models the genome with contiguous splice graphs and quantifies RNA-seq reads against these graphs. After read alignment, alternative splicing events are defined as a set of skipped nodes or connecting edges in the splice graph, and each alternative splicing event is quantified using expectation maximisation. The differential splicing events are then inferred by comparing the control and treatment conditions. Finally, Leafcutter identifies and quantifies novel and known alternative splicing events by focusing on intron excisions. Leafcutter analyses split-mapped short reads to locate overlapping introns that share splice sites. The detected introns are represented in a graph, of which the connected components form intron clusters (i.e., introns sharing donor or acceptor splice sites). For each cluster, alternative intron usage is quantified, and intron clusters with differentially excised introns are inferred.

SpliCeAT integrates these three tools and executes them using Snakemake to produce a set of high-confidence differential splicing events (Figure 1a), details of which can be found in Supplementary Figure 1. Briefly, RNA-seq read data from two experimental conditions are aligned to the reference genome using STAR two-pass alignment. MAJIQ, Whippet and Leafcutter are then executed concurrently to detect differential splicing events between the two conditions. The differential splicing events inferred by each tool are then filtered, keeping events produced by each tool with a junction percent spliced in (PSI) of |ΔPSI| ≥ 0.2 and either probability *p* ≥ 0.95 (for MAJIQ and Whippet) or FDR < 0.05 (for Leafcutter). Only events that are detected by two or more tools are considered high-confidence differential splicing events and retained for downstream use.

SpliCeAT also runs StringTie to produce a reference-guided transcriptome assembly (Figure 1a). Transcripts assembled by StringTie but not contained in the original reference transcriptome are used to augment the reference transcriptome only if they contain high-confidence differential splicing events as determined as described above.

### SpliCeAT detects high-confidence differential cryptic splicing events in embryonic mouse brain

SpliCeAT was run on short-read bulk RNA-seq data from the cortex and thalamus of a 14.5 day old embryonic mice with TDP-43 deleted from neural stem cells (cKO mice) and control mice (ctr mice). For the cortex, MAJIQ reported 172 genes with differential splicing, while Whippet and Leafcutter reported 837 and 221 genes respectively. Of these genes, only 96 genes were reported by all three tools to contain differential splicing (Figure 1b). Similarly, only 121 common genes were detected by all three tools from a total of 1024 unique splicing genes detected in the thalamus samples, demonstrating the low concordance of results between these tools. Taking splicing events reported by at least two tools, SpliCeAT reports a total of 215 and 266 high-confidence differential splicing events in the cortex and thalamus tissues, respectively (Figure 1b).

Various types of differential splicing events were detected, including cassette exons (included and skipped), retained introns, alternative splice sites, alternative first/last exon, alternative transcription start sites, and alternative polyadenylation sites. Of these types, cassette exons and alternative splice sites make up the majority of differential splicing events in the cortex and thalamus. In the cortex, 57.7 % (124/215) of differential splicing events were cassette exons, and 30.7 % (66/215) were alternative splice sites. Similarly, in the thalamus, cassette exons and alternative splice sites comprised 65.0 % (173/266) and 27.4 % (73/266) of differential splicing events respectively. Most detected cassette exons are novel inclusions of intronic regions (termed cryptic exons) that are otherwise unannotated and not seen in controls. ALS-related genes such as *Adnp2, Atxn2, Drosha, Camk1g, Synj2bp* and *Unc13a* were all detected to possess cryptic exon inclusions in cKO samples (Figure 1c and Supplementary Figure 2a). These cryptic exons coincide positionally with TDP-43 CLIP binding sites, indicating that the lack of TDP-43 protein in cKO state allowed otherwise repressed splice sites of cryptic exons to become active, thus leading to their inclusions. Furthermore, sequencing track examples of whole intron retention in *Atraid* and exon extension due to a novel splice acceptor in *Fam135a* are provided in Figure 1c.

Gene ontology enrichment was also performed for the set of genes detected to have differential splicing events in both cortex and thalamus samples. Nervous system processes such as synapse organisation, regulation of cell morphogenesis and regulation of neurotransmitters were significantly enriched among differential splicing genes across both tissues (Supplementary Figure 2b), suggesting that these nervous system processes may get dysregulated due to splicing defects.

### High-confidence differential splicing events are experimentally validated with long-read Iso-seq and RT-qPCR

Of the 215 high-confidence splicing events predicted by SpliCeAT for the embryonic mouse cortex samples, the transcript assembly segment of SpliCeAT was able to assemble 94.9 % of these events (204/215). SpliCeAT’s assembly of high-confidence splicing events is highly accurate, as about 91.6 % (197/215) of these high-confidence events were detected in Iso-seq transcripts (Figure 2a). Of the 18 high-confidence events from SpliCeAT not detected by Iso-seq, four events were randomly selected for validation by quantitative RT-PCR. We selectively amplified the novel exon-intron junctions from SpliCeAT and found that all four novel cryptic exons were experimentally validated (Figure 2b), suggesting that these are indeed true splicing events that were not picked up by Iso-Seq. This could likely be due to short cryptic exon lengths that were not captured by Iso-seq’s alignment tool or due to low long read sequencing depth. Nevertheless, the high-confidence differential splicing events detected and assembled into transcripts by SpliCeAT could indeed be experimentally validated by long-read Iso-seq.

**Figure 2.**
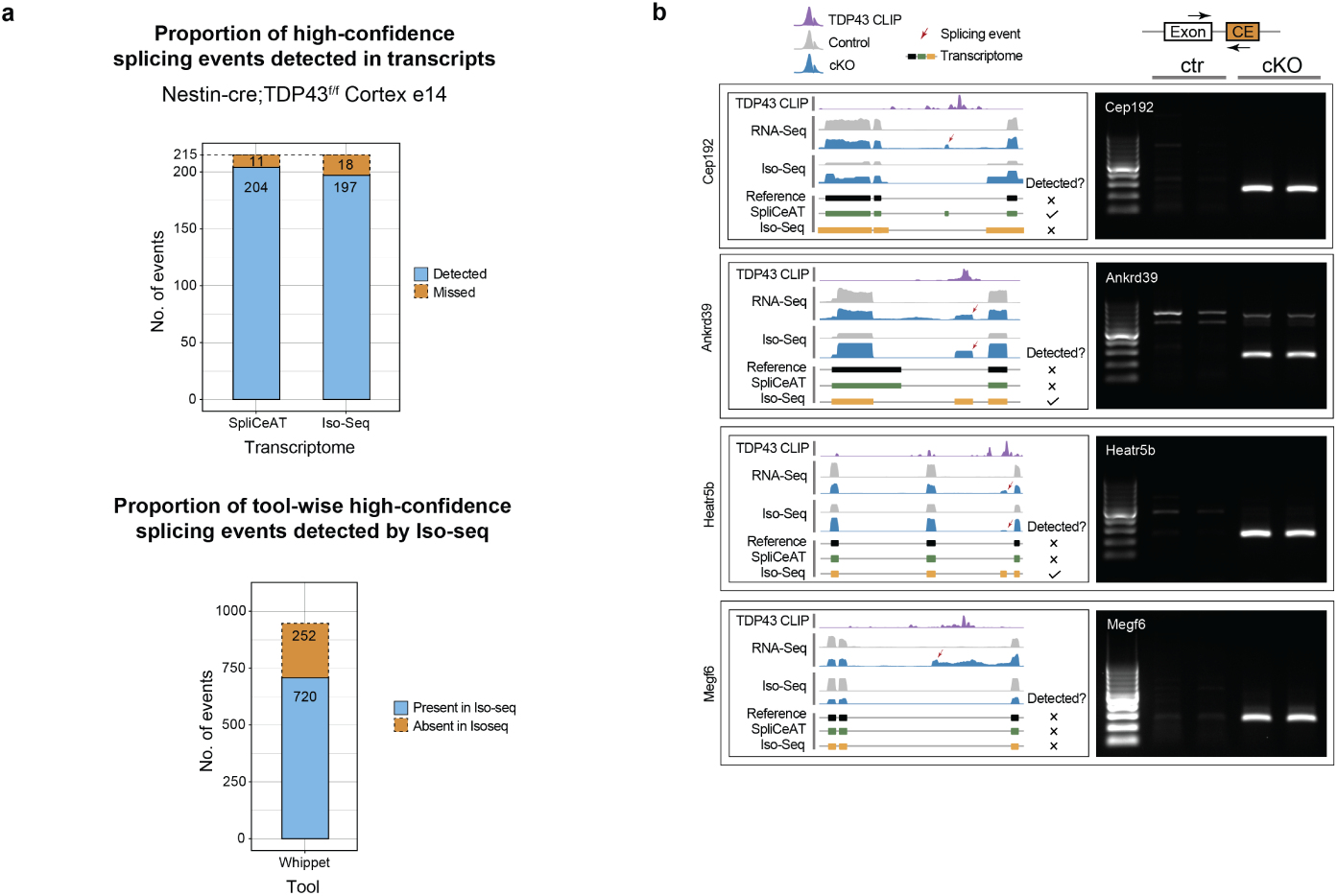
High Confidence Splicing events validated through Iso-Seq and RT-qPCR. (a) Proportion of high-confidence splicing events that were detected by SpliCeAT and Iso-seq in mouse cortex samples (top). Proportion of splicing events detected by Whippet that were also detected by Iso-seq (bottom). (b) Quantitative RT-PCR to detect cryptic exons in of four genes *Cep192*, *Ankrd39*, *Heatr5b* and *Megf6*.

Furthermore, novel splicing events from embryonic mice with TDP-43-depleted neural progenitor cells were compared with differential splicing events detected from TDP-43-negative neuronal nuclei from postmortem ALS-FTD patient brain samples [27]. Cryptic events in four genes (*Kalrn, Unc13a, Poldip3, Rnf144a*) were found to be conserved across both species (Supplementary Figure 2c), suggesting that aberrant splicing in these genes may play an important role in TDP-43-meditated pathogenesis. Indeed, all four genes have been shown to be associated with ALS [9, 28–31]. TDP-43-mediated aberrant splicing of UNC13A in humans has been reported to cause impaired neurotransmission, the rescue of which ameliorated the effect [9]. Additionally, UNC13A polymorphisms which cause aberrant splicing have been strongly associated with ALS risk [29] and anti-sense oligonucleotide-based therapeutics targeting aberrant UNC13A splicing are currently being considered [28]. Gene tracks of the cryptic splicing regions in these four genes in both mouse and human RNA-seq data are presented in Supplementary Figure 2d. The detected cryptic splicing events were also independently validated using RT-qPCR in mouse cortex samples, where ctr samples only contained one band while cKO samples separated into two bands, with the heavier band corresponding to the RNA section containing the cryptic exon. The inclusion of novel, and potentially disruptive, cryptic exons within these genes could likely lead to disruption of nervous system development in mouse embryos.

### SpliCeAT’s augmented transcriptome enables transcript-level analysis of novel splice events

Generation of an augmented transcriptome. When working with organisms that have a well characterised reference transcriptome, transcriptome-based RNA-seq quantification is often preferred to transcriptome assembly. SpliCeAT aims to retain the speed, accuracy, and consistency of transcriptome-based quantification while broadening its application to situations where a standard reference transcriptome is insufficient. It does this by augmenting the reference transcriptome (RefTx) file with assembled novel transcripts containing high-confidence differential splicing events. Details of the augmentation process are shown in Supplementary Figure 3. Briefly, StringTie is used for reference-guided transcript assembly of all RNA-seq samples. The resulting transcript assemblies (in GTF format) contain both reference transcripts and unannotated/novel transcripts, and these assemblies are merged across all samples to produce a set of unique, non-redundant transcripts. In order to avoid low-confidence or false positive novel transcripts assembled by StringTie, novel transcripts within this merged assembly undergo a filtering step to retain only transcripts that contain high-confidence differential splicing events as detected previously. Finally, these filtered novel transcripts containing high-confidence splicing events are added to the RefTx to construct the final augmented transcriptome (AugTx), which is used for downstream quantification and differential abundance analysis.

In our samples, reference-guided transcript assembly of control and TDP-43 cKO mice cortex samples using StringTie yielded a total of 154,930 transcripts, of which 12,696 were novel transcripts. Although StringTie has a higher novel transcript detection rate than other transcript assemblers like Cufflinks [32] and Bayesembler [33], it overestimates novel transcripts and has a low precision rate [34–36]. Indeed, out of the 12,696 novel transcripts assembled by StringTie, only 32.9 % (4,171/12,696) were experimentally validated and detected in long-read Iso-Seq. This reiterates the importance of filtering the assembled novel transcripts to only include those containing high-confidence differential splicing events. After filtering using the 215 high-confidence splicing events in cortex samples, we obtain total of 389 high-confidence novel transcripts from 170 genes. These novel transcripts were then added to the RefTx file to create the AugTx (Supplementary Figure 3).

Transcript-level analysis of novel splice events (E.g. *Adnp2*): As discussed previously, differential transcript expression analysis using the RefTx is unable to capture novel transcript expression, as novel transcripts are by default not accounted for in the RefTx. This can lead to a misrepresentation of isoform composition changes. We first illustrate this with the simple case of the *Adnp2* gene in our TDP-43 cKO mice (Figure 3). *Adnp2* has a single reference transcript Adnp2-201; in the cKO and expresses a cryptic cassette exon in the cKO mice, which was experimentally validated by Iso-seq (Figure 3a). The inclusion of the cryptic cassette exon leads to the assembly and addition of two novel transcripts: Adnp2-CE1 and Adnp2-CE2 in the AugTx (Figure 3a-b). Differential transcript expression analysis using the RefTx alone results in a significant upregulation of reference transcript Adnp2-201 (log_2_ FC = 0.72; *p*_adj_ = 0.001, Figure 3c). However, the differential expression results change drastically upon inclusion of the novel transcripts into the transcriptome (i.e. AugTx). The reference transcript now becomes significantly downregulated (log_2_ FC = −1.09; *p*_adj_ = 0.035) while the novel transcripts are significantly upregulated (Adnp2-CE1: log_2_ FC = 4.36; *p*_adj_ = 0.099, Adnp2-CE2: log_2_ FC = 1.68; *p*_adj_ = 0.021). This suggests that in the AugTx, reads mapping to the cryptic exon are now correctly assigned to the cryptic exon-containing transcripts, instead of being incorrectly mapped to the reference transcript Adnp2-201 as in the RefTx. Additionally, data from long-read Iso-seq corroborates the downregulation of the reference transcript Adnp2-201 (log_2_ FC = −3.82) and upregulation of cryptic exon-containing transcripts (log_2_ FC = 6.85, Figure 3c). This example in *Adnp2* underscores the relevance and importance of using an AugTx for accurate quantification of isoforms, particularly in cases where novel inclusions of the transcriptome are expected in samples.

**Figure 3.**
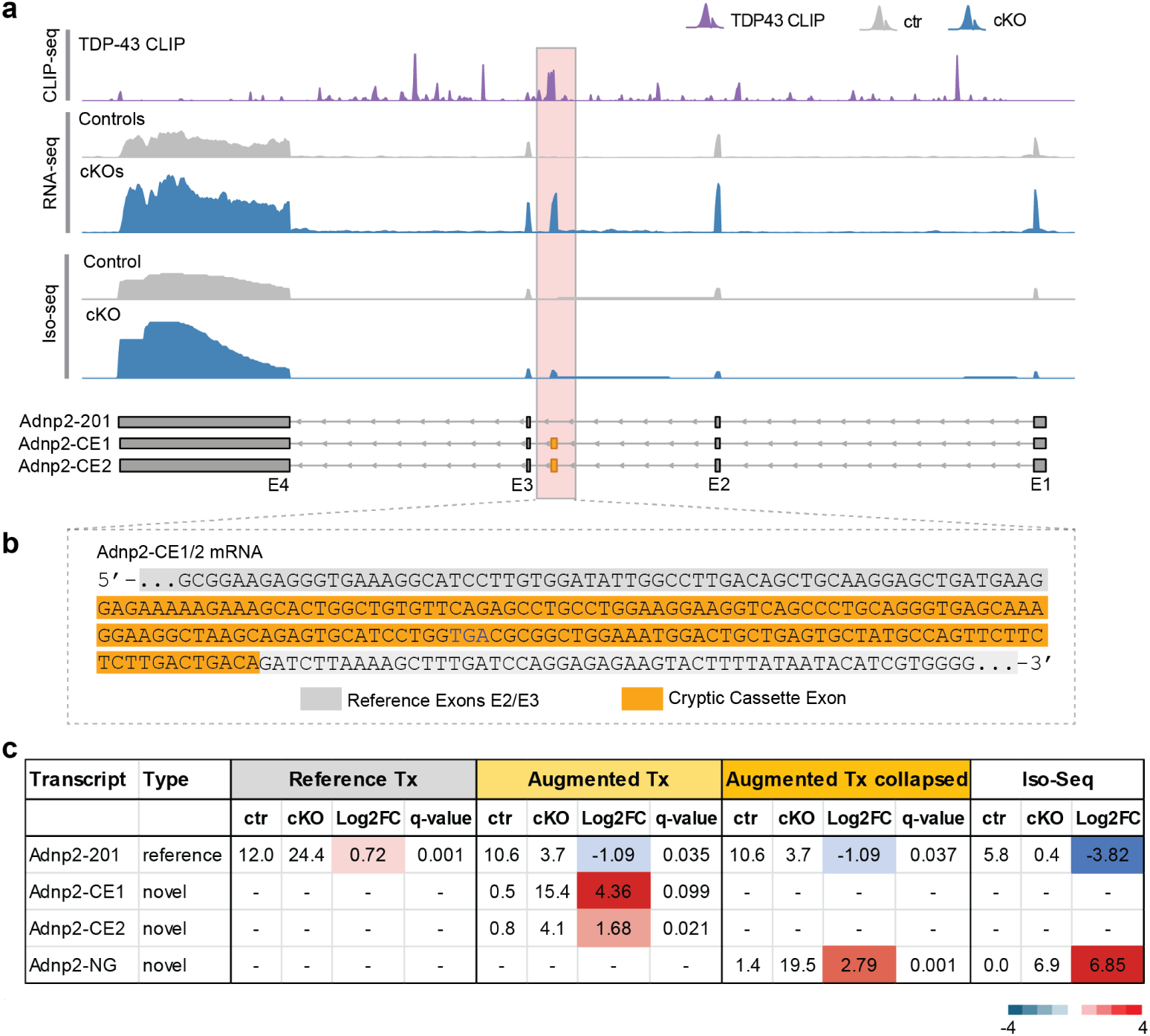
An augmented transcriptome allows for accurate quantification of *Adnp2*transcripts. (a) Visualisation of the *Adnp2* gene region with tracks from TDP-43 CLIP-seq binding sites, short read RNA-Seq reads from control and TDP-43 cKO mice, Iso-seq reads from ctr and cKO mice, followed by gene models containing the reference transcript and two novel transcripts incorporating the cryptic exon (highlighted in orange). (b) The sequence of *Adnp2* transcript region with the cryptic exon which incorporates a premature terminating codon in the open reading frame. (c) The mean transcripts per million (TPM) values in control and cKO mice and associated log2 fold change (log_2_ FC) and *p*_adj_for *Adnp2* transcripts using the reference transcriptome alone, augmented transcriptome (uncollapsed and collapsed) and from Iso-seq.

Collapsing novel transcripts using Lancaster aggregation: Although filtering for high-confidence novel transcripts is an imperative step, there may be cases where novel transcripts assembled by StringTie do not differ from each other substantially. For example, novel transcripts Adnp2-CE1 and Adnp2-CE2 only differ by 4 bp at the 5′-start exon (Figure 3a). Such small differences in exon boundaries, especially in the 5′ start exon or 3′ end exon, were observed across many novel transcripts containing splice events and were also observed in Iso-seq. This is attributed to one of the long-recognised problems of transcript assemblers, which is the inaccurate/fuzzy annotation of 5′ and 3′ ends of transcripts [37]. Additionally, for short-read data, it is often challenging to assign short reads spanning the cryptic region to just one transcript or another with high confidence, especially in genes with multiple long reference transcripts consisting of alternative exons. Furthermore, in such cases, StringTie assembles all possible exon permutations of cryptic exons into the reference transcripts, many of which may not reflect the actual underlying biology. In view of this, multiple novel transcripts with such small differences in exon boundaries can thus be treated as one isoform in order to simplify data interpretation. In order to do this, we collapsed transcripts containing the same high-confidence differential splicing event into one novel transcript cluster (termed as novel group; NG) using Lancaster aggregation of transcript adjusted *p*-values (*p*_adj_) implemented in Sleuth [38, 39]. Using this approach, Adnp2’s augmented gene model simplifies to one reference transcript Adnp2-201, which is downregulated in cKO mice (log_2_ FC = −1.09; *p*_adj_ = 0.037), and a novel transcript cluster Adnp2-NG, which is upregulated in cKO mice (log_2_ FC = 2.79; *p*_adj_ = 0.001, Figure 3c).

### An augmented transcriptome enables accurate quantification of global isoform composition changes

Next, we illustrate the importance of the AugTx on the level of the global transcriptome. Differential transcript expression analysis using RefTx yielded 511 differentially expressed transcripts (DETs) (*p*_adj_ < 0.05), while using the AugTx yielded 649 DETs (Figure 4d), with a radically distinct global differential transcript profile (Figure 4a-b). Not only are many novel transcripts such as Ube2d1-NG and Ppp6c-NG (marked in orange in Figure 4b) highly dysregulated, but less significant reference transcripts like Fam135a-201 and Abr-202 (in Figure 4a) now show up as highly dysregulated in the AugTx (in Figure 4b). This is due to a higher accuracy in transcript quantification using the AugTx, as reads mapping to differentially splicing events are now correctly assigned to novel transcripts, instead of being incorrectly attributed to reference transcripts or discarded. In Figure 4c, we observe the contrasts in transcript significance between using the Ref Tx and AugTx. AugTx captures the majority of the significant transcripts also reported by Ref Tx (465/511), with 184 additional transcripts (orange in colour) reported as significant (Figure 4d).

**Figure 4.**
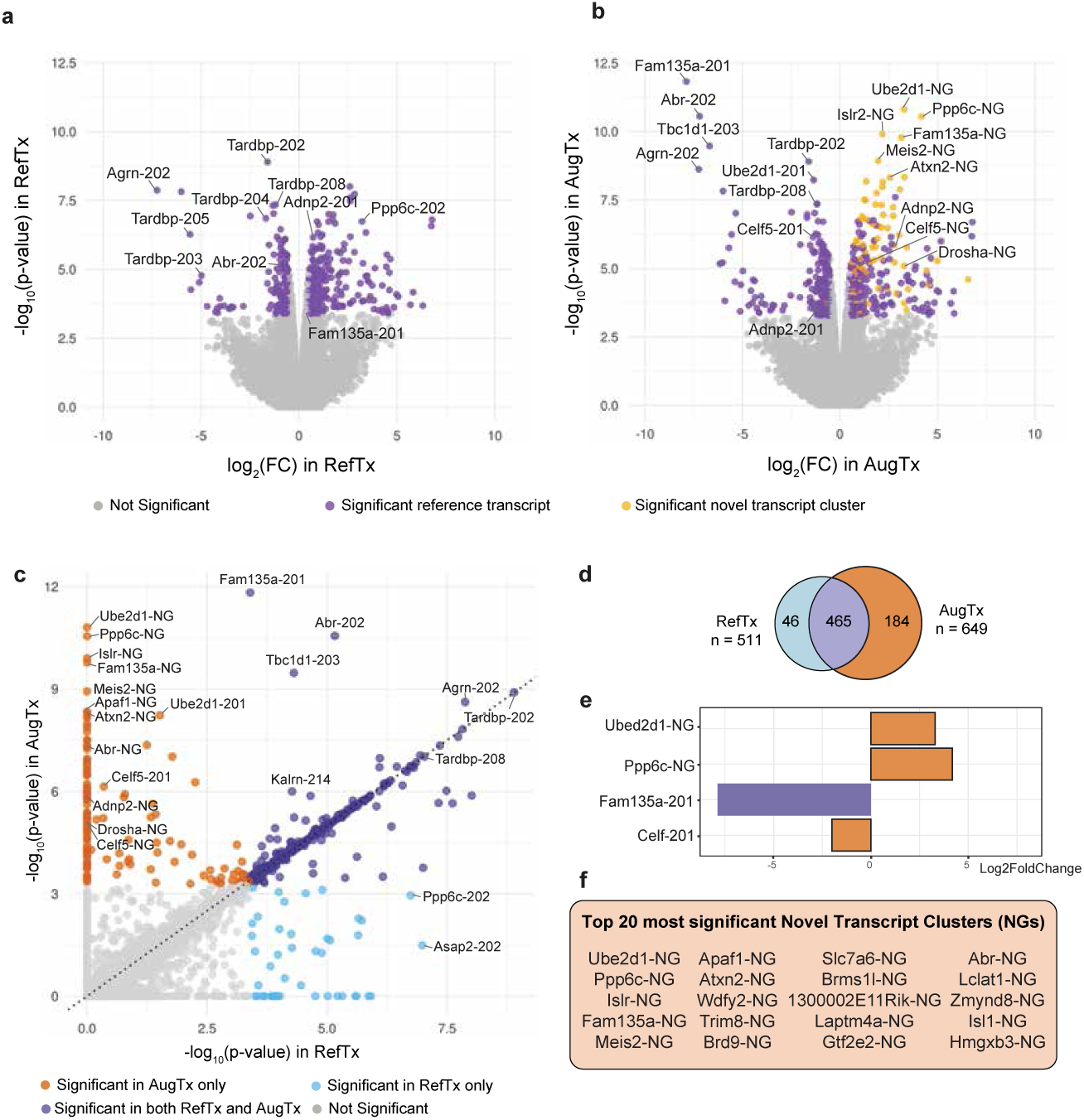
An augmented transcriptome enables accurate quantification of global isoform composition changes. (a) Volcano plot of differentially expressing transcripts in cKO mice using the RefTx model alone. (b) Volcano plot of differentially expressing transcripts in cKO mice using the collapsed AugTx model which includes novel transcripts with cryptic splicing events. Many new transcripts (highlighted in orange), which could not be captured by the RefTx, are now significantly differentially expressed. (c) Distribution of −log_10_ *p* values of differentially expressing transcripts using the RefTx versus the AugTx model. Transcripts in purple on the diagonal are significant using both models while transcripts in blue are significant only using the RefTx model. There are a large number of transcripts in orange which are quantified as significantly differentially expressing only using the AugTx. These include both reference transcripts and novel transcripts containing novel splicing events (NG transcripts). (d) Overlap between the significant differentially expressed transcripts between RefTx and AugTx. (e) log_2_ FC of the top few significantly differentially expressed transcripts in the AugTx. (f) Top 20 differentially expressed novel transcript clusters in the augmented transcriptome.

This suggests that in analyses of aberrant splicing conditions, solely using the publicly available RefTx is insufficient to account for the presence of novel transcripts generated due to treatment conditions, such as depletion of splicing regulator TDP-43 in our study. Thus, before performing differential analysis, it is important to first account for any novel transcripts that may be present in treatment samples. As presented above, this can be achieved by performing transcript assembly to generate novel transcripts, followed by augmenting them into the transcriptome to create an AugTx specific to experimental samples.

## Discussion

The human and mouse genomes have, not surprisingly, the most completely annotated transcriptomes of all vertebrates. This relatively complete annotation means that researchers can benefit from using the indexed transcript sequences directly when analysing RNA-sequencing experiments [38, 40, 41]. But those annotations do not capture the products of misregulation leading to the formation of aberrant transcripts and downstream functional loss in proteins.

Different splice event detection tools using a variety of approaches can detect novel splicing events, albeit with limited agreement. Our approach with SpliCeAT is to conservatively augment the reference transcriptome with assembled transcripts that are also confidently supported by detection of novel splice events from the data.

Using SpliCeAT we were able to identify high-confidence novel splicing events integrated from three different splicing detection tools. These high-confidence novel splicing events could be verified independently by long-read Iso-seq and RT-qPCR. With our pipeline, we found that, in the brain tissue of embryonic mice with TDP-43 depleted in neural stem cells, many ALS disease-associated genes such as *Adnp2, Atxn2, Drosha, Camk1g, Synj2bp* and *Unc13a* contained novel splicing events. This is consistent with literature, where cryptic exon incorporation in *UNC13A* transcripts has been demonstrated to be a key contributing factor of disease pathogenesis in ALS patients, with therapeutics targeting this mechanism currently being developed [9, 28, 29]. Furthermore, *ATXN2* trinucleotide expansions have also been shown to increase ALS risk [42], and intermediate expansions have been demonstrated to be causal in heritable ALS [43], with a phase I clinical trial targeting it underway [44, 45]. Our study is the first to report cryptic exon inclusions in *Atxn2* in mice models. Similarly, although *DROSHA* has been known to be implicated in neurodegeneration as seen in spinal muscular atrophy [46], we are the first to report TDP-43-mediated aberrant splicing as a potential mechanism for neurodegeneration.

A key part of the SpliCeAT pipeline is to focus on identifying high-confidence novel splicing events which can be validated experimentally. However, identifying high-confidence events and transcripts comes with the trade-off of a potentially high false negative rate as seen in Figure 2a. Here, 720 out of 972 novel splicing events predicted by Whippet could be observed in experimental Iso-seq data, but only 155 events from Whippet met our criteria for high-confidence events used in our analysis. In view of this, users can therefore exercise discrimination in choosing the stringency filters of high-confidence events as per their needs, with some users potentially preferring to use a full union of all splicing events predicted by the three tools, rather than just the union of all intersections between the tools, as presented in this paper. This flexibility is readily incorporated in the pipeline.

Our analyses with SpliCeAT demonstrate the importance of using an augmented transcriptome for accurate estimation of global transcriptomic changes, particularly in cases when one expects novel splicing events that are unlikely to be found in the reference. Another advantage of the augmented transcriptome is that it can be expanded to include more novel transcripts assembled across various published RNA-seq data to generate a sort of “meta”-augmented transcriptome. This expanded augmented transcriptome can be updated frequently with additional novel transcripts uncovered from new datasets or in different experimental conditions, providing a useful resource for the community to perform a quick scan for mis-splicing in their own datasets.

Furthermore, collapsing of novel transcripts using Lancaster *p*-value aggregation is of great advantage when one wants a quick overview of global isoform composition changes in their dataset, as most isoforms containing splicing events either vary at the 5′ or 3′ exons or with a small base pair difference in splice acceptor/donor sites. Additionally, in the case of complex splicing with multiple differential splicing events per gene, there is generally insufficient confidence in assembling the correct transcript with short-read RNA-seq data. In these cases, aggregating all novel transcripts into a novel transcript cluster is a sensible method to preserve correct transcript quantification without explicitly requiring the novel transcript models. In the SpliCeAT pipeline, users have the choice of whether or not to collapse novel splicing events.

## Conclusion

Splicing misregulation can result in aberrant transcripts and is implicated in various diseases. Although current methods have enabled detection of aberrant splicing events, they are usually limited to the splice events themselves. Here, we present a novel pipeline that not only detects and quantifies novel splicing, but also augments the canonical transcriptome with novel transcripts containing these novel splice events. This enables accurate quantification of novel transcripts in the presence of aberrant splicing and improved estimation of global isoform composition changes. SpliCeAT’s accuracy is validated using RT-qPCR and long-read Iso-seq, making it a valuable and accurate tool for the splicing research community.

## Methods

### Mouse models

The study was carried out under protocols approved by the Institutional Animal Care and Use Committee (IACUC) of the National University of Singapore and was in compliance with Association for Assessment of Laboratory Animal Care (AAALAC) guidelines for animal use. The mice were maintained at C57BL/6J background and housed in groups with individually ventilated cages under a 12:12-hour light/dark cycle and access to food/water *ad libitum*. Conditional TDP-43 (Tardbp-fl/fl) mice (stock number 017591) and Nestin-Cre (stock number 003771) [47] were purchased from the Jackson Laboratory.

### RNA-sequencing and data processing

RNA extracted from the cortex and thalamus of 14.5-day-old developing mouse embryos of genotype Tardbp-fl/fl (control mice) and Nestin-Cre;Tardbp-fl/fl mice (cKO mice) (n = 4 each) were used to prepare RNA-seq libraries using the TruSeq Stranded mRNA Library Prep Kit (Illumina). Illumina HiSeq4000 was used to generate 150 bp-long paired-end reads and yielded 30-40 million reads per sample.

FASTQ files were aligned to mouse reference genome (GRCm39) with STAR (v. 2.7.9a) with the following modified parameters: –outSAMunmapped Within –twopassMode Basic –sjdbOverhang 149. Mouse reference genome and gene annotation files (vM29) were obtained from GENCODE. The resulting aligned BAM files were sorted and indexed with samtools (v. 1.13).

### SpliCeAT pipeline

Differential splicing detection: SpliCeAT combines the following splicing detection tools using a Snakemake pipeline: MAJIQ/Voila (v. 2.4.dev3+g85d07819), Whippet (v. 1.6.1) and Leafcutter (v. 0.2.9), for the purpose of identifying high-confidence differential splicing events and novel cryptic exons present in the data.

For MAJIQ/Voila, mouse gene annotation file (GENCODE vM29 primary assembly GFF3 file) was used to build a transcriptome index in the form of gene-wise splice graphs. Novel splice junctions were added to the index by supplementing RNA-seq read information from the sample aligned BAM files. Reads from sample BAM files were then quantified against the index to identify differential splicing events. MAJIQ was run with the default parameters. Voila was used as an interactive server to display and obtain the list of differential splicing events.

For Whippet, a transcriptome index was built using the mouse gene annotation file (GENCODE vM29 primary assembly GTF file) with the following modified parameters: –bammin-reads 20. The index was then supplemented with merged TDP-43 cKO BAM files for inclusion of novel splice junctions. Sample FASTQ files were aligned and quantified against the transcriptome index under default parameters to obtain a list of differential splicing events.

For detection of splicing events by Leafcutter, all exon-exon junctions with minimum split-read anchor length of 8 bp were extracted from sample BAM files with regtools (v. 0.5.2) junction extract under the following parameters: -a 8 -m 50 -M 500000. Introns corresponding to exon-exon junctions were clustered together into intron clusters using leafcutter cluster.py with the following modified parameter: -l 500000. Differential testing of intron clusters was carried out using leafcutter ds.py with the following modified parameters: -i 1 -g 3, to obtain a list of differential intron clusters.

High-confidence differential splicing events with the following criteria were selected: MAJIQ (|ΔPSI| ≥ 0.2; *p* ≥ 0.95), Whippet (|ΔPSI| ≥ 0.2; *p* ≥ 0.95), and Leafcutter (|ΔPSI| ≥ 0.2; FDR < 0.05). Differential splicing events identified by at least two tools (i.e., union of all intersects) were retained and used for downstream creation of the augmented transcriptome.

Generation of an augmented transcriptome: Reference-guided (using GENCODE mouse vM29 primary assembly GTF file) transcript assembly was performed for each sample BAM file using StringTie (v. 2.2.1) with -j 20. All resulting transcript assemblies were merged to produce a set of unique, non-redundant transcripts in GTF format. All novel assembled transcripts were extracted from the assembled GTF and converted into a GRanges object in R using GenomicRanges (v. 1.50.2), and only transcripts that contain the high-confidence differential splicing events detected previously were retained. Any remaining novel assembled transcripts that did not contain these high-confidence differential splicing events were discarded.

Gffread (v. 0.12.7) was used to obtain the ribonucleotide sequence (in FASTA format) of the novel transcripts containing high-confidence differential splicing events. The FASTA file of novel transcripts was then concatenated with the mouse reference transcriptome (GENCODE GRCm39 FASTA file) to create the augmented transcriptome.

### Quantification and differential analysis of transcripts

kallisto (v. 0.46.1) index was used to generate a transcriptome index from the augmented transcriptome described previously using default parameters. kallisto quant was used to quantify transcripts present within samples, and sleuth (v. 0.30.1) was used for differential transcript expression analysis, differential transcript utilisation analysis and differential gene expression analysis. In order to perform differential transcript expression analysis with transcript groups instead of individual transcripts, novel transcripts containing the same cryptic exon within the same gene were collapsed into a single novel transcript group. The transcript count of the novel transcript group was calculated as the sum of transcript per million counts of all its constituent transcripts within the same group. Significance of the collapsed transcript groups was calculated using Lancaster *p*-value aggregation in sleuth. The transcript-to-gene mapping used in sleuth was similarly augmented with the high-confidence novel transcripts. All RNA-seq BAM alignments were visualised on Gviz (v. 1.42.0) in R.

### RT-qPCR

Total RNA was extracted from cortex using Trizol reagent. After DNase treatment using RQ1 RNase-Free water, 1 microgram of RNA was reverse transcribed using Maxima First Strand cDNA Synthesis Kit for RT-qPCR (Thermo Fisher). mRNA levels were determined using Maxima SYBR Green RT-qPCR master mix (Thermo Fisher). The expression levels of the gene of interests were normalised to the extended ΔCT, obtained from three reference genes, e.g., RPL13, RPLP0, and ARHGDIA. Primers for the selected mouse genes are as follows in Table 1.

**Table 1.**
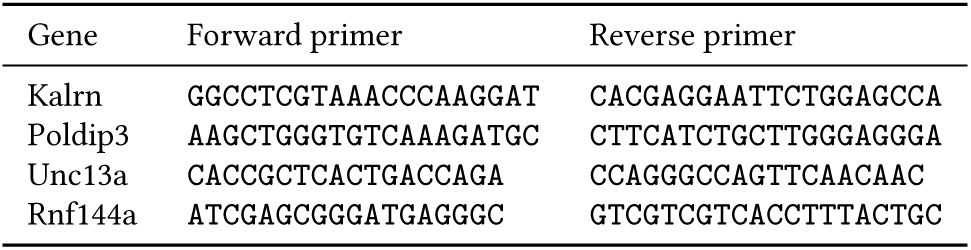
List of Primers.

### Whole transcriptome Iso-seq library preparation, SMRT sequencing and data processing

Iso-seq was performed on RNA from cortex samples of control (n = 1) and cKO (n = 3) 14.5 day old mouse embryos. Long read Iso-seq library preparation was carried out by Novogene. Briefly, first strand cDNA was synthesised from polyA+ tail using NEBNext’s reverse transcriptase. Template switching, extension and amplification were performed to generate double-stranded cDNA, which were then ligated with universal hairpin adapters. After magnetic bead purification, sequencing primers were annealed to SMRTbell templates, followed by binding of the sequence polymerase to annealed templates. Libraries were pooled and sequenced on PacBio Sequel II/Ile system. The circularised DNA template was sequenced with multiple passes to generate subreads. Consensus was called from subreads to produce highly accurate long reads (HiFi reads).

QC and analysis of HiFi reads was performed using the Isoseq3 pipeline (v. 3.8.2, https://github.com/PacificBiosciences/IsoSeq). Primers were removed from HiFi reads using lima (v. 2.7.1) in specialised Iso-seq mode to generate full-length reads. Isoseq3 refine was then used to remove concatemer HiFi reads and perform polyA tail trimming to produce full-length, non-concatemer reads. These reads were then clustered using isoseq3 cluster under default parameters to generate full-length transcripts, which were subsequently mapped to mouse reference genome (GENCODE vM29) using pbmm2 (v. 1.7.0) under ISOSEQ preset to obtain high-quality (predicted accuracy ≥99 %) aligned transcripts. The aligned transcripts were collapsed using isoseq3 collapse based on genomic coordinates to produce collapsed transcript models in GFF format.

### Characterisation of Iso-seq isoforms

Collapsed Iso-seq isoforms in GFF format were characterised using SQANTI3 (v. 5.1.1, https://github.com/ConesaLab/SQANTI3) using GENCODE vM29 mouse gene annotation GTF and reference genome. Reference CAGE peaks for GRCm39 mouse genome, polyA motif list for mouse and human and full-length read counts information for each isoform were also utilised for characterisation. During characterisation, each isoform was assigned to one of the following structural classifications when compared to reference transcripts in the gene annotation: full splice match, incomplete splice match, novel in catalog, novel not in catalog, antisense, genic intron, genic genomic and intergenic.

### Rarefaction analysis

Cupcake (v. 29.0.0, https://github.com/Magdoll/cDNA_Cupcake) was used to perform rarefaction curve analysis. Collapsed isoform GFF, FASTQ and abundance files were used to perform subsampling with a minimum full-length read count of 2 and step size 1000. Rarefaction curves at the gene, isoform and transcript category levels were plotted in R.

### Detection of differential splicing events in Iso-seq and RNA-seq transcripts

High-confidence differential splicing events were converted into coordinate ranges in BED files using bedtools (v. 2.30.0). Iso-seq GFF file and StringTie GTF file were converted into GRanges objects and the ranges were subset to ‘exon’ types only. The subset GRanges objects were then converted into BED files containing coordinate ranges. Bedtools intersect with -f 1 was used to obtain the intersection of the differential splicing events BED file and the Iso-seq/StringTie exon BED files to determine the number of differential splicing events detected by Iso-seq and StringTie respectively.

In order to ascertain the accuracy of StringTie’s transcript assembly process, Iso-seq and StringTie transcript models were first subset to include only transcripts containing high-confidence novel cryptic exons, and gffcompare (v. 0.12.8) was subsequently used to compare Iso-seq and StringTie transcript models.

For all Iso-seq analysis, Iso-seq TPM transcript expression values were calculated as follows: 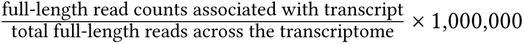. Only multi-exonic transcripts were considered.

## Declarations

### Funding

This work was supported by grants from the Swee Liew-Wadsworth Endowment fund to S.-C. Ling and I. Agrawal; MOE Tier1/NUHS seed grant National University of Singapore (NUS), National Medical Research Council (OFIRG23jul-0050), Singapore and Ministry of Education (MOE-T2EP30220-0014), Singapore to S.-C. Ling; and MOE Tier 1 grant funding to G. Tucker-Kellogg.

### Competing interests

The authors declare no competing interests.

### Ethics approval

The study was carried out under protocols approved by the Institutional Animal Care and Use Committee (IACUC) of the National University of Singapore and was in compliance with Association for Assessment of Laboratory Animal Care (AAALAC) guidelines for animal use.

### Consent to participate

All studies were carried out under protocols approved by the Institutional Animal Care and Use Committee (IACUC) of the National University of Singapore, and were in compliance with Association for Assessment of Laboratory Animal Care (AAALAC) guidelines for animal use.

### Consent for publication

Not applicable.

### Availability of data and materials

The RNA-Seq datasets used in the paper are available at GSE288457.

### Code availability

The code used in this study is available at: https://github.com/GTK-lab/SpliCeAT.

### Authors’ contributions

Authors’ contribution: GTK and SCL conceived and coordinated the study. YSL, IA, SCL and GTK wrote the paper. IA, YSL, ES, WYH, SCL and GTK designed, performed and analysed the experiments shown in Figures 1–4 and Supplemental Figures 1-3. All authors reviewed the results and approved the final version of the manuscript.

## Acknowledgements

We thank both the Tucker-Kellogg and Ling laboratory members for their support, discussion, and suggestions.

## Notes

https://www.ncbi.nlm.nih.gov/geo/query/acc.cgi?acc=GSE288457

### Competing Interest Statement

The authors have declared no competing interest.

## References

[1] Su CH, D D, Tarn WY. Alternative Splicing in Neurogenesis and Brain Development. Frontiers in Molecular Biosciences. 2018;5. Tex.ids= suAlternativeSplicingNeurogenesis2018a publisher: Frontiers.

[2] Raj B, Blencowe BJ. Alternative Splicing in the Mammalian Nervous System: Recent Insights into Mechanisms and Functional Roles. Neuron. 2015 Jul;87(1):14–27. Publisher: Elsevier. 10.1016/j.neuron.2015.05.004.

[3] Fagnani M, Barash Y, Ip JY, Misquitta C, Pan Q, Saltzman AL, et al. Functional coordination of alternative splicing in the mammalian central nervous system. Genome Biology. 2007 Jun;8(6):R108. 10.1186/gb-2007-8-6-r108.

[4] Frankish A, Diekhans M, Jungreis I, Lagarde J, Loveland J, Mudge JM, et al. GENCODE 2021. Nucleic Acids Research. 2020 Dec;49(D1):D916–D923. 10.1093/nar/gkaa1087.

[5] Noguchi S, Arakawa T, Fukuda S, Furuno M, Hasegawa A, Hori F, et al. FANTOM5 CAGE profiles of human and mouse samples. Scientific Data. 2017 Aug;4(1):170112. Publisher: Nature Publishing Group. 10.1038/sdata.2017.112.

[6] Prudencio M, Belzil VV, Batra R, Ross CA, Gendron TF, Pregent LJ, et al. Distinct brain transcriptome profiles in C9orf72-associated and sporadic ALS. Nature Neuroscience. 2015 Aug;18(8):1175–1182. Number: 8 Publisher: Nature Publishing Group. 10.1038/nn.4065.

[7] Ling JP, Pletnikova O, Troncoso JC, Wong PC. TDP-43 repression of nonconserved cryptic exons is compromised in ALS-FTD. Science. 2015 Aug;349(6248):650–655. 10.1126/science.aab0983.

[8] Mehta PR, Brown AL, Ward ME, Fratta P. The era of cryptic exons: implications for ALS-FTD. Molecular Neurodegeneration. 2023 Mar;18(1):16. 10.1186/s13024-023-00608-5.

[9] Ma XR, Prudencio M, Koike Y, Vatsavayai SC, Kim G, Harbinski F, et al. TDP-43 represses cryptic exon inclusion in the FTD-ALS gene UNC13A. Nature. 2022 Mar;603(7899):124–130. 10.1038/s41586-022-04424-7.

[10] Seddighi S, Qi YA, Brown AL, Wilkins OG, Bereda C, Belair C, et al. Mis-spliced transcripts generate de novo proteins in TDP-43-related ALS/FTD. bioRxiv: The Preprint Server for Biology. 2023 Jan;p. 2023.01.23.525149. 10.1101/2023.01.23.525149.

[11] Melamed Z, López-Erauskin J, Baughn MW, Zhang O, Drenner K, Sun Y, et al. Premature polyadenylation-mediated loss of stathmin-2 is a hallmark of TDP-43-dependent neurodegeneration. Nature Neuroscience. 2019 Feb;22(2):180–190. 10.1038/s41593-018-0293-z.

[12] Baughn MW, Melamed Z, López-Erauskin J, Beccari MS, Ling K, Zuberi A, et al. Mechanism of STMN2 cryptic splice-polyadenylation and its correction for TDP-43 proteinopathies. Science (New York, NY). 2023 Mar;379(6637):1140–1149. 10.1126/science.abq5622.

[13] Sun M, Bell W, LaClair KD, Ling JP, Han H, Kageyama Y, et al. Cryptic Exon Incorporation Occurs in Alzheimer’s Brain lacking TDP-43 Inclusion but Exhibiting Nuclear Clearance of TDP-43. Acta neuropathologica. 2017 Jun;133(6):923. 10.1007/s00401-017-1701-2.

[14] Estades Ayuso V, Pickles S, Todd T, Yue M, Jansen-West K, Song Y, et al. TDP-43-regulated cryptic RNAs accumulate in Alzheimer’s disease brains. Molecular Neurodegeneration. 2023 Aug;18(1):57. 10.1186/s13024-023-00646-z.

[15] Teboul R, Grabias M, Zucman-Rossi J, Letouzé E. Discovering cryptic splice mutations in cancers via a deep neural network framework. NAR Cancer. 2023 Mar;5(2):zcad014. 10.1093/narcan/zcad014.

[16] Li YI, Knowles DA, Humphrey J, Barbeira AN, Dickinson SP, Im HK, et al. Annotation-free quantification of RNA splicing using LeafCutter. Nature Genetics. 2018 Jan;50(1):151–158. 10.1038/s41588-017-0004-9.

[17] Sterne-Weiler T, Weatheritt RJ, Best AJ, Ha KCH, Blencowe BJ. Efficient and Accurate Quantitative Profiling of Alternative Splicing Patterns of Any Complexity on a Laptop. Molecular Cell. 2018 Oct;72(1):187–200.e6. Publisher: Elsevier. 10.1016/j.molcel.2018.08.018.

[18] Vaquero-Garcia J, Barrera A, Gazzara MR, González-Vallinas J, Lahens NF, Hogenesch JB, et al. A new view of transcriptome complexity and regulation through the lens of local splicing variations. eLife. 2016 Feb;5:e11752. Publisher: eLife Sciences Publications, Ltd. 10.7554/eLife.11752.

[19] Kahles A, Ong CS, Zhong Y, Rätsch G. SplAdder: identification, quantification and testing of alternative splicing events from RNA-Seq data. Bioinformatics (Oxford, England). 2016 Jun;32(12):1840–1847. 10.1093/bioinformatics/btw076.

[20] Middleton R, Gao D, Thomas A, Singh B, Au A, Wong JJL, et al. IRFinder: assessing the impact of intron retention on mammalian gene expression. Genome Biology. 2017 Mar;18(1):51. 10.1186/s13059-017-1184-4.

[21] Romero JP, Muniategui A, De Miguel FJ, Aramburu A, Montuenga L, Pio R, et al. EventPointer: an effective identification of alternative splicing events using junction arrays. BMC genomics. 2016 Jun;17:467. 10.1186/s12864-016-2816-x.

[22] Mancini E, Rabinovich A, Iserte J, Yanovsky M, Chernomoretz A. ASpli: integrative analysis of splicing landscapes through RNA-Seq assays. Bioinformatics (Oxford, England). 2021 Sep;37(17):2609–2616. 10.1093/bioinformatics/btab141.

[23] Denti L, Rizzi R, Beretta S, Vedova GD, Previtali M, Bonizzoni P. ASGAL: aligning RNA-Seq data to a splicing graph to detect novel alternative splicing events. BMC bioinformatics. 2018 Nov;19(1):444. 10.1186/s12859-018-2436-3.

[24] Mehmood A, Laiho A, Venäläinen MS, McGlinchey AJ, Wang N, Elo LL. Systematic evaluation of differential splicing tools for RNA-seq studies. Briefings in Bioinformatics. 2020 Dec;21(6):2052–2065. 10.1093/bib/bbz126.

[25] Trincado JL, Entizne JC, Hysenaj G, Singh B, Skalic M, Elliott DJ, et al. SUPPA2: fast, accurate, and uncertainty-aware differential splicing analysis across multiple conditions. Genome Biology. 2018 Mar;19(1):40. Tex.ids= trincadoSUPPA2FastAccurate2018a, trincadoSUPPA2FastAccurate2018b, trincadoSUPPA2FastAccurate2018c. 10.1186/s13059-018-1417-1.

[26] Fenn A, Tsoy O, Faro T, Rößler FM, Dietrich A, Kersting J, et al. Alternative splicing analysis benchmark with DICAST. NAR Genomics and Bioinformatics. 2023 Jun;5(2):lqad044. 10.1093/nargab/lqad044.

[27] Liu EY, Russ J, Cali CP, Phan JM, Amlie-Wolf A, Lee EB. Loss of Nuclear TDP-43 Is Associated with Decondensation of LINE Retrotransposons. Cell reports. 2019 Apr;27(5):1409–1421.e6. 10.1016/j.celrep.2019.04.003.

[28] Willemse SW, Harley P, van Eijk RPA, Demaegd KC, Zelina P, Pasterkamp RJ, et al. UNC13A in amyotrophic lateral sclerosis: from genetic association to therapeutic target. Journal of Neurology, Neurosurgery, and Psychiatry. 2023 Aug;94(8):649–656. 10.1136/jnnp-2022-330504.

[29] Brown AL, Wilkins OG, Keuss MJ, Hill SE, Zanovello M, Lee WC, et al. TDP-43 loss and ALS-risk SNPs drive mis-splicing and depletion of UNC13A. Nature. 2022 Mar;603(7899):131–137. 10.1038/s41586-022-04436-3.

[30] Gittings LM, Alsop EB, Antone J, Singer M, Whitsett TG, Sattler R, et al. Cryptic exon detection and transcriptomic changes revealed in single-nuclei RNA sequencing of C9ORF72 patients spanning the ALS-FTD spectrum. Acta Neuropathologica. 2023 Sep;146(3):433–450. 10.1007/s00401-023-02599-5.

[31] Shiga A, Ishihara T, Miyashita A, Kuwabara M, Kato T, Watanabe N, et al. Alteration of POLDIP3 Splicing Associated with Loss of Function of TDP-43 in Tissues Affected with ALS. PLoS ONE. 2012 Aug;7(8):e43120. 10.1371/journal.pone.0043120.

[32] Trapnell C, Williams BA, Pertea G, Mortazavi A, Kwan G, van Baren MJ, et al. Transcript assembly and quantification by RNA-Seq reveals unannotated transcripts and isoform switching during cell differentiation. Nature Biotechnology. 2010 May;28(5):511–515. Publisher: Nature Publishing Group. 10.1038/nbt.1621.

[33] Maretty L, Sibbesen JA, Krogh A. Bayesian transcriptome assembly. Genome Biology. 2014;15(10):501. 10.1186/s13059-014-0501-4.

[34] Shao M, Kingsford C. Accurate assembly of transcripts through phase-preserving graph decomposition. Nature Biotechnology. 2017 Dec;35(12):1167–1169. 10.1038/nbt.4020.

[35] Kovaka S, Zimin AV, Pertea GM, Razaghi R, Salzberg SL, Pertea M. Transcriptome assembly from long-read RNA-seq alignments with StringTie2. Genome Biology. 2019 Dec;20(1):278. 10.1186/s13059-019-1910-1.

[36] Voshall A, Moriyama EN. Next-Generation Transcriptome Assembly: Strategies and Performance Analysis. In: Abdurakhmonov IY, editor. Bioinformatics in the Era of Post Genomics and Big Data. IntechOpen; 2018. p. 15–36. Tex.ids= voshallNextGeneration-TranscriptomeAssembly2018a. Available from: https://www.intechopen.com/chapters/59634.

[37] Steijger T, Abril JF, Engström PG, Kokocinski F, RGASP Consortium, Hubbard TJ, et al. Assessment of transcript reconstruction methods for RNA-seq. Nature Methods. 2013 Dec;10(12):1177–1184. 10.1038/nmeth.2714.

[38] Pimentel H, Bray NL, Puente S, Melsted P, Pachter L. Differential analysis of RNA-seq incorporating quantification uncertainty. Nature Methods. 2017 Jul;14(7):687–690. Number: 7 Publisher: Nature Publishing Group. 10.1038/nmeth.4324.

[39] Yi L, Pimentel H, Bray NL, Pachter L. Gene-level differential analysis at transcript-level resolution. Genome Biology. 2018 Apr;19(1):53. 10.1186/s13059-018-1419-z.

[40] Bray NL, Pimentel H, Melsted P, Pachter L. Near-optimal probabilistic RNA-seq quantification. Nature Biotechnology. 2016 May;34(5):525–527. Number: 5 Publisher: Nature Publishing Group. 10.1038/nbt.3519.

[41] Patro R, Duggal G, Love MI, Irizarry RA, Kingsford C. Salmon provides fast and bias-aware quantification of transcript expression. Nature Methods. 2017 Apr;14(4):417–419. 10.1038/nmeth.4197.

[42] Elden AC, Kim HJ, Hart MP, Chen-Plotkin AS, Johnson BS, Fang X, et al. Ataxin-2 intermediate-length polyglutamine expansions are associated with increased risk for ALS. Nature. 2010 Aug;466(7310):1069–1075. 10.1038/nature09320.

[43] Lee T, Li YR, Ingre C, Weber M, Grehl T, Gredal O, et al. Ataxin-2 intermediate-length polyglutamine expansions in European ALS patients. Human Molecular Genetics. 2011 May;20(9):1697–1700. 10.1093/hmg/ddr045.

[44] Amado DA, Davidson BL. Gene therapy for ALS: A review. Molecular Therapy: The Journal of the American Society of Gene Therapy. 2021 Dec;29(12):3345–3358. 10.1016/j.ymthe.2021.04.008.

[45] Biogen. A Phase 1/2 Multiple-Ascending-Dose Study With a Long-Term Open-Label Extension to Assess the Safety, Tolerability, Pharmacokinetics, Pharmacodynamics, and Effect on Disease Progression of BIIB105 Administered Intrathecally to Adults With Amyotrophic Lateral Sclerosis With or Without Poly-CAG Expansion in the ATXN2 Gene. clinicaltrials.gov; 2023. NCT04494256. Submitted: July 30, 2020. Available from: https://clinicaltrials.gov/study/NCT04494256.

[46] Gonçalves IdCG, Brecht J, Thelen MP, Rehorst WA, Peters M, Lee HJ, et al. Neuronal activity regulates DROSHA via autophagy in spinal muscular atrophy. Scientific Reports. 2018 May;8(1):7907. 10.1038/s41598-018-26347-y.

[47] Tronche F, Kellendonk C, Kretz O, Gass P, Anlag K, Orban PC, et al. Disruption of the glucocorticoid receptor gene in the nervous system results in reduced anxiety. Nature Genetics. 1999 Sep;23(1):99–103. Number: 1 Publisher: Nature Publishing Group. 10.1038/12703.

