## Supplementary Figures for "Integrated detection and quantification of aberrant transcripts with novel splicing events"

### Supplementary material for: Integrated detection and quantification of aberrant transcripts with novel splicing events*

February 3, 2025

* Agrawal, Lim, Sanford, Ho, Ling, and Tucker-Kellogg

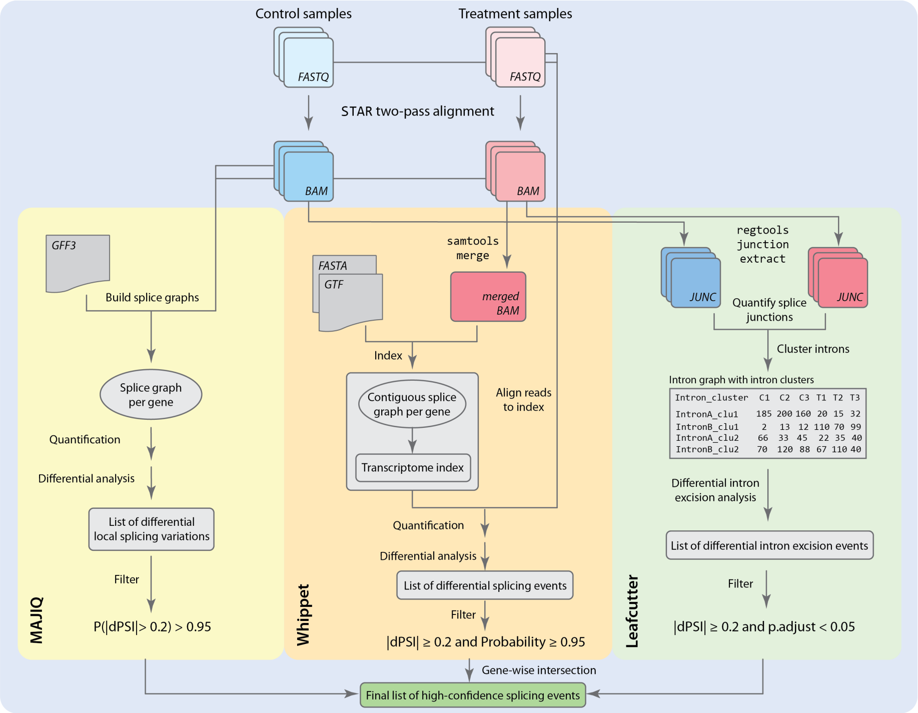


Supplementary Figure 1: **Overview of high confidence splicing events in SpliCeAT.** RNA-seq FASTQ files of two conditions are aligned to the reference genome using STAR two-pass alignment in order to generate indexed BAM alignment files. MAJIQ, Whippet and Leafcutter are then performed separately to detect differential splicing events between the two conditions. Presented here are the broad steps taken within each software to detect differential splicing events. To ensure that only high-confidence differential splicing events are retained, SpliCeAT filters all differential splicing events returned by each tool with the following filter: $|\Delta PSI| \geq 0.2$ and $P \geq 0.95$ (for MAJIQ and Whippet) or FDR $< 0.05$ (for Leafcutter). Out of these filtered events, only those identified by at least two software are retained and used for downstream creation of the augmented transcriptome.

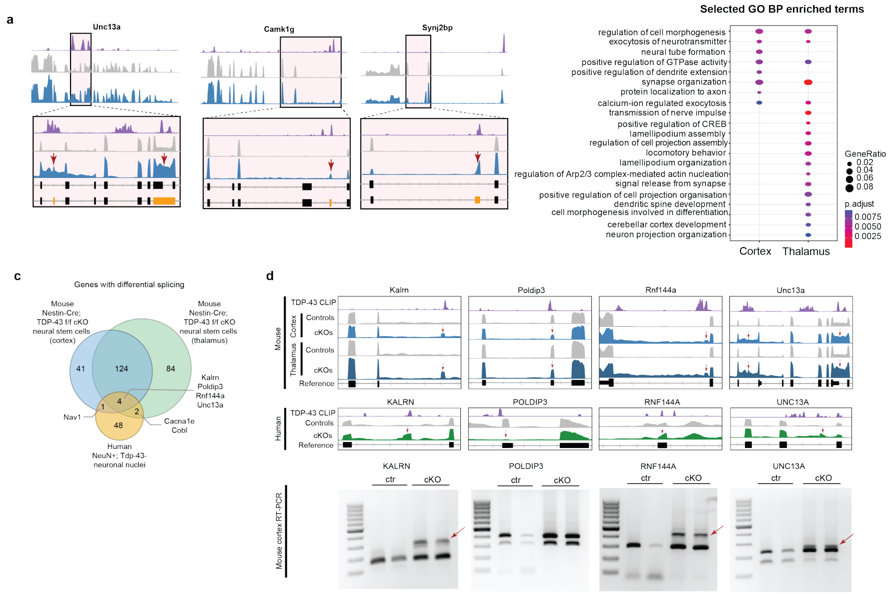


Supplementary Figure 2: **Widespread differential splicing detected in Nestin-Cre; TDP-43 f/f conditional knockout embryonic mouse brain and its similarities to human TDP-43 negative neuronal nuclei**. (a) Alignment of RNA-seq read coverage tracks for TDP-43 CLIP (purple peaks), TDP-43 f/f controls (grey peaks) and Nestin-Cre; TDP-43 f/f cKOs (blue peaks) for selected ALS-related genes detected to contain differential splicing events in cortex and thalamus. Read coverage tracks plotted here are of thalamus tissue, although similar peak patterns are found in cortex as well. Black gene models represent transcript models belonging to the reference transcriptome, while orange bars represent novel exons in the augmented transcriptome. (b) Enriched gene ontology biological processes in the genes containing differential splicing events of the TDP-43 cKO cortex and thalamus tissues. (c) Venn diagram showing genes detected to contain differential splicing events in the TDP-43 cKO mouse cortex (blue), TDP-43 cKO mouse thalamus (green), and human TDP-43 negative neuronal nuclei (yellow). Overlapping genes found between datasets are labelled. Kalrn, Poldip3, Rnf144a and Unc13a were found to contain differential splicing events in all three datasets. (d) RNA-seq alignment tracks for the four overlapping genes detected to have differential splicing events in TDP-43 mouse cortex/thalamus and human TDP-43 negative neuronal nuclei datasets. Red arrows on the alignment tracks indicate the location of either a cryptic exon or exon skipping. RT-PCR in mouse cortex validates the presence of cryptic exons (indicated by red arrows) detected in Kalrn, Rnf144a and Unc13a. Primers were designed to flank the detected cryptic exons.

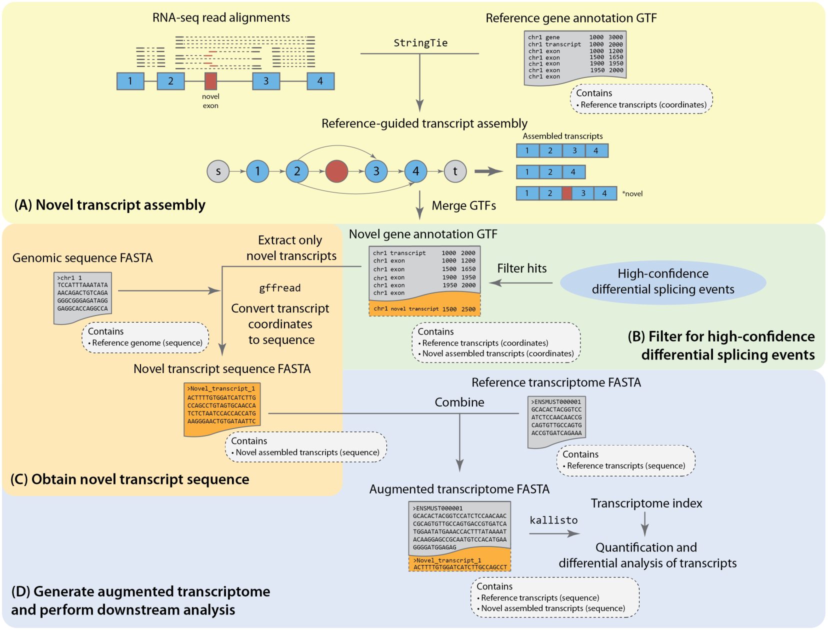


Supplementary Figure 3: **Overview of augmented transcriptome construction in SpliCeAT**. Reference-guided transcript assembly is performed for each sample BAM file using StringTie. All assemblies were merged to produce a set of unique, non-redundant transcripts. All novel assembled transcripts were extracted from the assembled GTF, and only transcripts that contain the high-confidence differential splicing events were retained. Gffread was used to obtain the nucleotide sequence (in FASTA format) of the novel transcripts containing high-confidence differential splicing events. The FASTA file of novel transcripts was concatenated with the reference transcriptome to create the augmented transcriptome.
